# Heart Rate n-Variability (HRnV): A Novel Representation of Beat-to-Beat Variation in Electrocardiogram

**DOI:** 10.1101/449504

**Authors:** Nan Liu, Dagang Guo, Zhi Xiong Koh, Andrew Fu Wah Ho, Marcus Eng Hock Ong

## Abstract

Heart rate variability (HRV) is a widely adopted tool for evaluating changes in cardiac autonomic regulation. The majority of efforts have focused on developing methods to assess HRV by deriving sophisticated parameters with linear and nonlinear techniques and adopting advanced signal processing tools for efficient noise removal and accurate QRS detection. In this paper, we propose a novel representation of beat-to-beat variation in an electrocardiogram (ECG), called heart rate n-variability (HRnV), as an alternative to conventional HRV measures. We derived two novel HRnV measures based on non-overlapped and overlapped RR intervals. We also conducted a simulation study by using an ECG record from the MIT-BIH Normal Sinus Rhythm Database to demonstrate the feasibility of calculating HRnV parameters. Among the time domain parameters, we observed that the values were generally incremental with the increase in *n*. We observed the same trend of changes for the frequency domain parameters. In the nonlinear analysis, the differences between HRV and HRnV from Poincare plot measures were obvious, while those from entropy and detrended fluctuation analysis metrics were not. HRnV measures enable us to augment conventional HRV measures with additional parameters. Although issues remain to be addressed regarding HRnV, we hope to stimulate a new stream of research on this new representation of HRV. HRnV is an important addition to HRV and will contribute to extending the landscape of current studies on HRV.

## 1 Introduction

Heart rate variability (HRV), a widely adopted tool for evaluating changes in cardiac autonomic regulation, is believed to be strongly associated with the autonomic nervous system. Due to its popularity in many clinical applications, the guidelines for HRV measurement, physiological interpretation and clinical use were published in 1996 [1]. Acharya et al. [2] presented a comprehensive review of the analytical methods used to measure HRV and applications of HRV. More recently, Billman [3] reviewed HRV from a historical perspective.

The aim of HRV analysis is to explore the beat-to-beat variation in an electrocardiogram (ECG). Over the years, numerous quantitative techniques have been adopted, improved, and implemented to analyze ECGs to capture these variations [4]. For example, geometrical methods are used to extract time domain parameters, Fourier transform analysis is implemented to derive frequency domain parameters, and detrended fluctuation analysis (DFA) is adopted to calculate nonlinear parameters.

HRV has gained a reputation for use in a broad range of clinical applications, particularly in cardiovascular research, in which reduced HRV is regarded as a significant predictor of adverse outcomes [5]. However, the impact of the autonomic nervous system on HRV remains controversial [3], leaving room for further clinical studies exploring novel engineered parameters to model beat-to-beat variation. So far, the majority of efforts have focused on deriving sophisticated parameters with linear and nonlinear techniques. Furthermore, researchers have focused on developing advanced signal processing tools for efficient noise removal and accurate QRS detection prior to calculating HRV parameters.

In this paper, we revisit RR intervals and, the foundations for computing HRV parameters, and propose heart rate n-variability (HRnV), a novel representation of beat-to-beat variation in ECGs. We have developed two specific HRnV measures as alternatives to conventional HRV measures and evaluated the feasibility of computing these new parameters. We will also discuss the merits, issues, and potential applications of HRnV measures, and suggest directions for future development.

## 2 Proposed Heart Rate n-Variability

We elaborate two measures of the novel HRnV representation, namely, HR*_n_*V and HR*_n_*V_m_. We will introduce the definitions of both of these measures and illustrate the differences between them and conventional HRV measures.

### 2.1 HR*_n_*V: A Novel Measure with Non-Overlapped RR Intervals

Prior to introducing the new HR*_n_*V measure, we define a new type of RR interval (RRI) called RR*_n_*I, where 1 ≤ *n* ≤ *N*, and 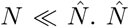 is the total number of RR intervals. When *n* =1, RR*_n_*I becomes conventional RRI. The definition of RR*_n_*I is illustrated in Fig. 1. Note that RR_1_I is equal to RRI. When *n* > 1, every *n* adjacent RRI is connected to form a new sequence of RR*_n_*Is. By using this strategy, we are able to create a maximum number of (*N* − 1) new RR*_n_*I sequences from the conventional single RRI sequence.

**Fig. 1.**
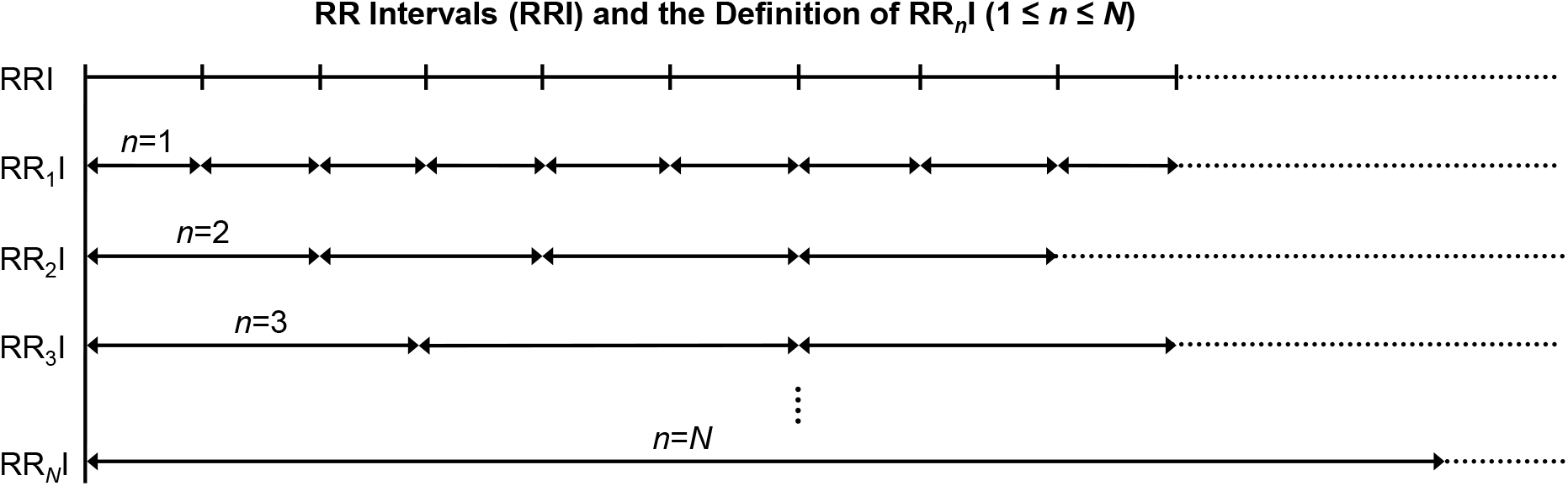
Illustration of RR intervals and the definition of RR*_n_*I where 1 ≤ *n* ≤ *N* and 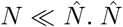 is the total number of RR intervals.

With these newly generated RR*_n_*I sequences, the calculation of HR*_n_*V parameters is straight-forward and can be accomplished by applying existing quantitative methods including time domain analysis, frequency domain analysis, and nonlinear analysis [1],[2]. The computed HR*_n_*V parameters include but are not limited to the following: the average of RR*_n_*Is (aRR*_n_*), standard deviation of RR*_n_*Is (sdRR*_n_*), square root of the mean squared differences between RR*_n_*Is (RMSSD*_n_*), the number of times that the absolute difference between 2 successive RR*_n_*Is exceeds 50 ms (NN50*_n_*), NN50*_n_* divided by the total number of RR*_n_*Is (pNN50*_n_*), the integral of the RR*_n_*I histogram divided by the height of the histogram (HR*_n_*V triangular index), low frequency (LF*_n_*) power, high frequency (HF*_n_*) power, approximate entropy (ApEn*_n_*), sample entropy (SampEn*_n_*), and detrended fluctuation analysis (DFA*_n_*), among others. We use the subscript n to indicate that the parameters are calculated from RR*_n_*I sequences.

As noted in the above description, HR*_n_*V is a novel measure based on newly generated, non-overlapped RR*_n_*Is. Next, we will introduce another novel measure, HR*_n_*V*_m_*, which is based on overlapped RRIs.

### 2.2 HR*_n_*V*_m_*: A Novel Measure with Overlapped RR Intervals

Like RR*_n_*I, which is used in HR*_n_*V to define the HR*_n_*V*_m_* measure, we introduce another type of RRI called RR*_n_*I*_m_* where 1 ≤ *n* ≤ *N*, 1 ≤ *m* ≤ *N* − 1, and 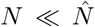. In the RR*_n_*I*_m_* sequence, *m* is used to indicate the level of overlapping between consecutive RR*_n_*I*_m_* sequences. As depicted in Fig. 2, (*n* − *m*) RRIs form the overlapped portions. Apparently, when *m* = *n*, RR*_n_*I*_m_* becomes RR*_n_*I. Therefore, the upper limit of *m* is *N* − 1. By controlling the overlap among newly generated RR*_n_*I*_m_* sequences, we are able to create a maximum number of (*N* × (*N* − 1)/2) RR*_n_*I*_m_* sequences (excluding the RR*_n_*I sequence) from the conventional single RRI sequence.

**Fig. 2.**
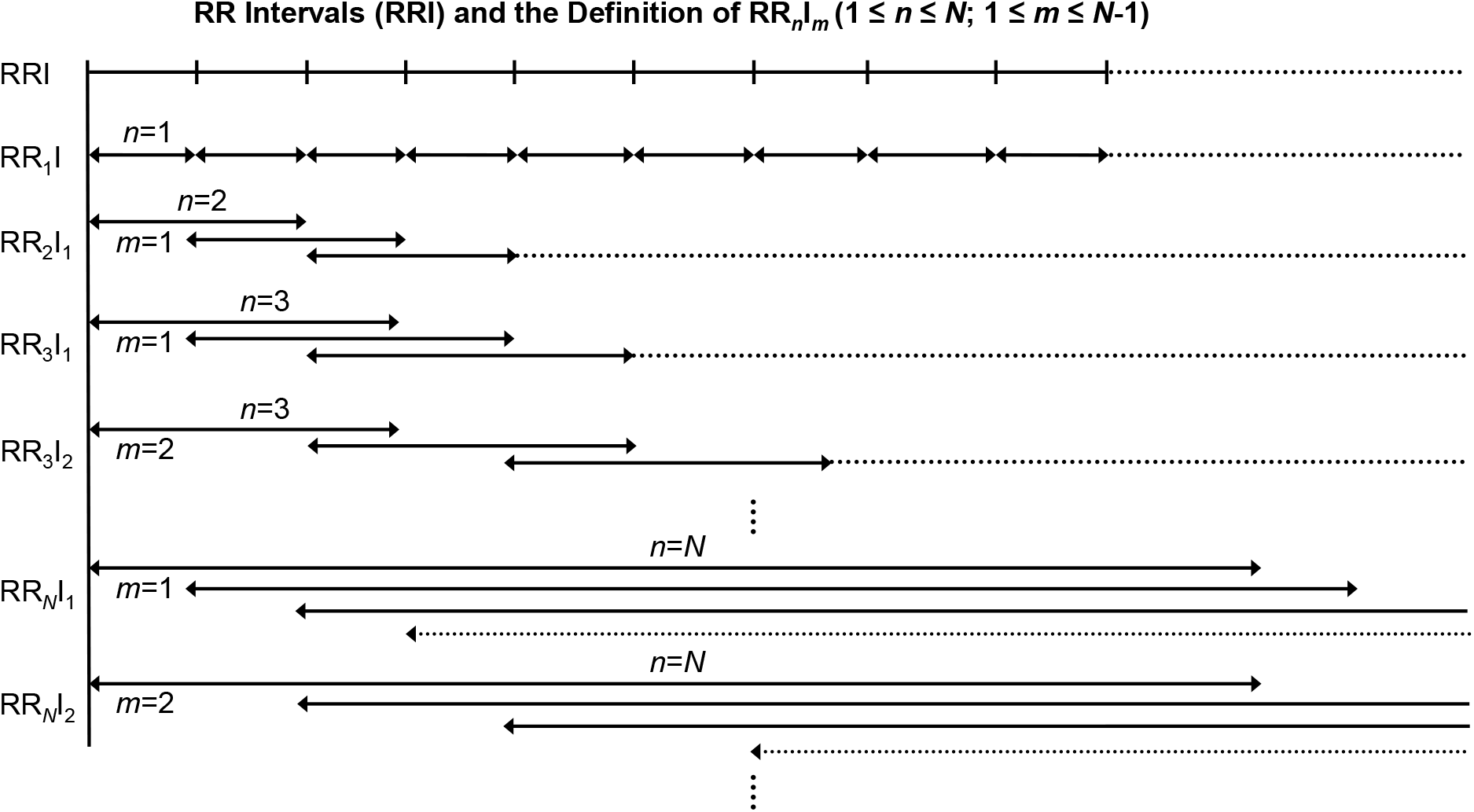
Illustration of RR intervals and the definition of RR*_n_*I*_m_* where 1 ≤ *n* ≤ *N*, 1 ≤ *m* ≤ *N* − 1, and 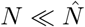.

For each of the newly created RR*_n_*I*_m_* sequences, we can apply time domain analysis, frequency domain analysis, and nonlinear analysis, to calculate HR*_n_*V*_m_* parameters. We add the superscript m to denote that the parameters are computed from RR*_n_*I*_m_* sequences. For example, the average RR*_n_*I*_m_* intervals and the sample entropy are written as 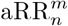 and 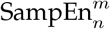, respectively.

The HR*_n_*V*_m_* measure extracts more information from the original RRI sequence than HR*_n_*V, by adopting a strategy of controlling sequence overlap. The HR*_n_*V*_m_* measure is particularly useful and suitable when ECG segments are short, and thus there are a limited number of RRIs.

## 3 Experiments

To evaluate the feasibility of calculating HRnV parameters, we conducted a simulation study by using the ECG record of subject #16265 from the MIT-BIH Normal Sinus Rhythm Database [6]. We applied the conventional Pan-Tompkins QRS detection algorithm including a bandpass filter (5-15 Hz), derivative filter, and moving average to detect QRS peaks. Subsequently, we extracted a 30-minute segment and derived the RR*_n_*I and RR*_n_*I*_m_* sequences from the original RRIs, where *n* ≤ 3. The conventional RRI, RR*_n_*I, and RR*_n_*I*_m_* sequences are illustrated in Fig. 3. We observed that there were no obvious changes in the waveforms of conventional and new RRIs. However, toward the end of sequences, we noted a spike in the original RRI but smoother areas in the RR*_n_*I and RR*_n_*I*_m_* sequences, indicating that sudden significant changes in adjacent R peaks could have been suppressed in the new RR*_n_*I and RR*_n_*I*_m_* representations in which multiple intervals were connected.

Based on the six RRI sequences shown in Fig. 3, we calculated HRV, HR*_n_*V, and HR*_n_*V*_m_* parameters (Table 1). Among the time domain parameters, we observed that the values were generally incremental with an increase in *n*. Special attention needs to be given to NN50 and pNN50, where 50 ms is the threshold to assess the difference between pairs of successive RRIs. Notably, in HRnV measure, the lengths of RR*_n_*I and RR*_n_*I*_m_* have been extended; thus, the threshold needs to be adjusted accordingly. As shown in Table 1, we used 50 ms as the default threshold for all calculations, since we did not aim to study specific parameters in this paper.

**Fig. 3.**
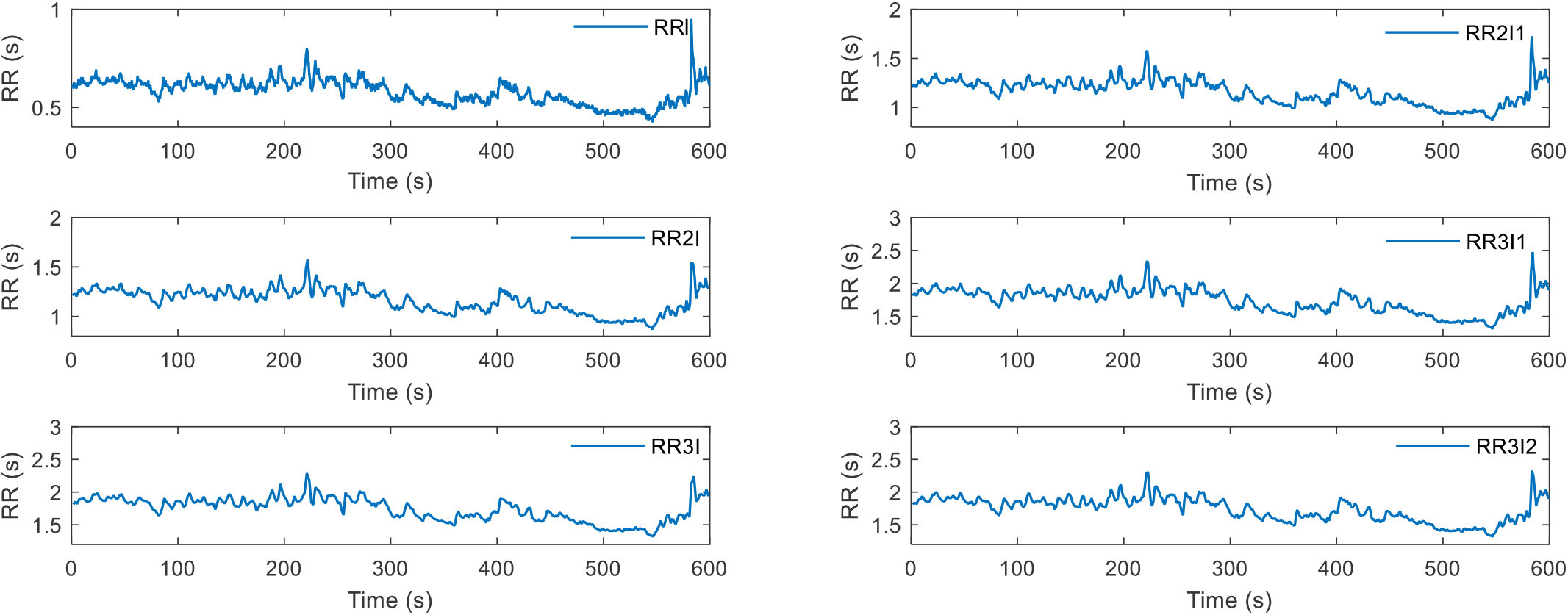
RR interval and its variations used for calculating HRnV parameters. The six RR interval sequences were RRI, RR2I, RR31, RR2I1, RR3I1, and RR3I2.

**TABLE 1.** Selected time domain, frequency domain, and nonlinear HRV and HRnV parameters based on a 30-minute ECG segment that was obtained from subject #16265 of the MIT-BIH Normal Sinus Rhythm Database.

| Parameters | HRV | HR<sub>2</sub>V | HR<sub>3</sub>V | HR<sub>2</sub>V<sub>1</sub> | HR<sub>3</sub>V<sub>1</sub> | HR<sub>3</sub>V<sub>2</sub> |
| --- | --- | --- | --- | --- | --- | --- |
| **Time Domain** |  |  |  |  |  |  |
| aRR (ms) | 651.2 | 1302.3 | 1953.4 | 1302.3 | 1953.5 | 1953.4 |
| sdRR (ms) | 81.0 | 160.0 | 237.8 | 159.9 | 237.7 | 237.7 |
| RMSSD (ms) | 25.9 | 58.9 | 99.1 | 34.8 | 41.3 | 73.5 |
| NN50 | 93 | 347 | 397 | 250 | 372 | 444 |
| pNN50 (%) | 3.4 | 25.1 | 43.2 | 9.1 | 13.5 | 32.2 |
| Triangular index | 16.8 | 35.4 | 35.4 | 38.4 | 48.4 | 49.3 |
| **Frequency Domain** |  |  |  |  |  |  |
| LF power (ms<sup>2</sup>) | 697.4 | 2648.5 | 5549.3 | 2669.1 | 5534.0 | 5581.8 |
| HF power (ms<sup>2</sup>) | 264.4 | 702.8 | 771.9 | 766.8 | 1017.5 | 940.4 |
| LF power norm (nu) | 72.5 | 79 | 87.8 | 77.7 | 84.5 | 85.6 |
| HF power norm (nu) | 27.5 | 21.0 | 12.2 | 22.3 | 15.5 | 14.4 |
| LF/HF | 2.638 | 3.769 | 7.190 | 3.481 | 5.439 | 5.935 |
| **Nonlinear** |  |  |  |  |  |  |
| Poincaré plot SD1 (ms) | 18.3 | 41.7 | 70.1 | 24.6 | 29.2 | 52.0 |
| Poincaré plot SD2 (ms) | 113.1 | 222.4 | 329.0 | 224.8 | 334.9 | 332.2 |
| Approximate entropy | 0.926 | 1.114 | 1.147 | 0.763 | 0.656 | 1.011 |
| Sample entropy | 0.860 | 1.028 | 1.049 | 0.699 | 0.614 | 0.930 |
| DFA, $\alpha_1$ | 1.270 | 1.167 | 1.089 | 1.426 | 1.594 | 1.252 |
| DFA, $\alpha_2$ | 0.988 | 0.923 | 1.027 | 1.003 | 1.023 | 0.938 |

Like the time domain parameters, we observed the same trend of changes in frequency domain parameters. The exception was HF power norm, in which the HRnV parameters were smaller than the HRV parameters. We also noticed that the change in LF power norm was marginal compared to the change in HF power norm, which resulted in a significant difference in LF/HF values between HRV and HRnV parameters. In nonlinear analysis, the differences between HRV and HRnV parameters obtained from Poincaré plot measures were obvious, while those obtained from entropy and DFA metrics were not. The experimental results reported above were for demonstration purposes; thus, they were not meant to provide physiological interpretations. Furthermore, we have to consider many factors, such as subject characteristics and the length of ECG records, in rigorous clinical studies to conduct in-depth investigations on HRnV parameters and their clinical uses.

## 4 Discussion and Future Directions

In this paper, we introduced HRnV, a novel representation of beat-to-beat variation in ECGs. We proposed two measures, namely, HR*_n_*V and HR*_n_*V*_m_*. HR*_n_*V is calculated based on non-overlapped RR*_n_*I sequences, while HR*_n_*V*_m_* is computed from overlapping RR*_n_*I*_m_* sequences. HRnV is not proposed to replace the conventional HRV; instead, this measure is a natural extension of HRV. HR*_n_*V and HR*_n_*V*_m_* measures enable us to create additional alternative parameters from raw ECGs, and thus empower the extraction of supplementary information. Therefore, HRnV is complementary to HRV because this measure represents additional beat-to-beat variations in an ECG.

We have witnessed plentiful clinical investigations using conventional HRV parameters in cardiology [7], diabetes [8], critical care [9], psychiatry [10], cancer [11], and so forth. Similarly, we foresee broad application opportunities for HRnV. With the augmented RR*_n_*I and RR*_n_*I*_m_* sequences, HRnV parameters could possibly capture more dynamic pattern changes in various aspects than HRV does.

Given the richness of HRnV parameters, there are many ways of applying these parameters for research and applications. We briefly categorize three approaches for the application of HRnV parameters:

1. Use individual HRnV measures as alternatives to the conventional HRV parameters.
2. Stack various HRnV measures to form a high dimensional feature vector for predictive modeling and disease associations.
3. Aggregate various HRnV measures to create an ensemble of different models [12] that are built upon individual HRnV measures.

Approaches 2) and 3) are particularly suitable for artificial intelligence and machine learning tools [13], where many methods are available for statistical decision making [14], variable selection [15], and data mining [16].

Although HRnV has promising capabilities for augmenting conventional HRV parameters, there are still many issues to address. First, HRnV lacks physiological interpretations of its numerous parameters. Second, the choice of parameters n and m is arbitrary, which has a considerable impact in various conditions. For example, HRnV may not be feasible for very short ECGs in which the number of RRIs is limited. Third, the calculation of certain HRnV parameters needs to be carefully evaluated and rigorously investigated. NN50 in conventional HRV is defined as the number of successive RRI pairs that differ by more than 50 ms. However, in HR*_n_*V and HR*_n_*V*_m_*, 50 ms seems to no longer be a valid indicator. Thus, is *n* × 50 ms or another value a reasonable number? Addressing these issues requires collaborative endeavors between clinician scientists and biomedical engineering researchers.

## 5 Conclusions

We proposed using multiple RRIs (with or without overlaps) to create novel HRnV measures to represent beat-to-beat variation. We illustrated the definitions of HR*_n_*V and HR*_n_*V*_m_* and evaluated the feasibility of parameter calculation. HRnV measures enable us to augment the conventional HRV measures with many more parameters. We have also discussed three approaches with which new HRnV parameters can be used and adopted to assist with existing research. Although some issues remain to be addressed, we hope to stimulate a new stream of research on HRnV, a novel representation of beat-to-beat variation in ECG. We believe that future endeavors in this field will lead to the possibility of in-depth evaluation of the associations between HRnV measures and various human diseases.

## Author Contributions

N. Liu conceived the idea of heart rate n-variability (HRnV), developed the HR*_n_*V and HR*_n_*V*_m_* measures, and wrote the first draft of the manuscript. N. Liu, D. Guo, and Z.X. Koh performed the experiments. All authors contributed to the evaluation of the HRnV measures and revision of the manuscript.

